# Subventricular zone adult mouse neural stem cells and human glioblastoma stem cells require insulin receptor for self-renewal

**DOI:** 10.1101/2020.03.10.985598

**Authors:** Shravanthi Chidambaram, Deborah Rothbard, Kaivalya Deshpande, Yvelanda Cajuste, Kristin M. Snyder, Eduardo Fajardo, Andras Fiser, Nikos Tapinos, Steven W. Levison, Teresa L. Wood

## Abstract

The insulin receptor (IR) is an evolutionarily conserved signaling protein that regulates development and cellular metabolism. IR signaling regulates neurogenesis in *Drosophila*; however, a specific role for the IR in maintaining adult neural stem cells (NSCs) in mammals has not been investigated. We show that conditionally deleting the IR reduces adult NSCs of the subventricular zone by ∼70% accompanied by a corresponding increase in progenitors. IR deletion produced hyposmia due to aberrant olfactory bulb neurogenesis. Interestingly, hippocampal neurogenesis was not perturbed nor were hippocampal dependent behaviors. Highly aggressive proneural and mesenchymal glioblastomas (GBMs) had high IR/insulin-like growth factor (IGF) pathway gene expression, and isolated glioma stem cells had an aberrantly high ratio of IR:IGF1R receptors. Moreover, IR knockdown inhibited proneural and mesenchymal GBM tumorsphere growth. Altogether, these data demonstrate that the IR is essential for a subset of normal NSCs as well as for brain tumor cancer stem cell self-renewal.

## Introduction

Insulin or insulin-like signaling is a highly conserved signaling system that regulates development as well as key cellular functions including glucose and lipid metabolism. Whereas *Drosophila* insulin-like peptides signal through a single IR, mammals possess several insulin- related signaling receptors that include the well-known metabolic IR-B isoform as well as the A splice variant of the IR (lacking exon 11 containing 12 amino acids), and the insulin-like growth factor type 1 receptor (IGF-1R). Unlike IR-B, the IR-A and IGF-1R regulate cell proliferation and have been reported to regulate stem cell self-renewal or progenitor amplification in various cell populations. The family of ligands for these receptors includes insulin, IGF-I and IGF-II with varying affinities for the different receptors (Belfiore et al., 2017; Boucher et al., 2014). At physiological levels, insulin binds with high affinity only to the two IR isoforms. Conversely, IGF-I binds with high affinity only to the IGF-1R. IGF-II, however, is unique among the ligands in that it binds with high affinity to the IGF-1R but also has high affinity for IR-A (Frasca et al., 1999). The function of IGF-II through IR-A has been the focus of several investigations, which demonstrate that this signaling loop is: 1) essential for embryonic development (Efstratiadis, 1998; Louvi et al., 1997), 2) elevated in certain types of tumors (Belfiore, 2007; Frasca et al., 1999), and 3) necessary in vitro for self-renewal of normal NSCs (Ziegler et al., 2014; Ziegler et al., 2015; Ziegler et al., 2012) and thyroid tumor stem cells (Malaguarnera et al., 2012).

Our recent studies demonstrated that IGF-II is an essential niche factor for several populations of adult stem cells, including the NSCs of the subventricular zone (SVZ) and the subgranular zone (SGZ) (Ziegler et al., 2019). In the adult central nervous system, IGF-II is predominantly produced by the choroid plexus, although it also is expressed at lower levels by hippocampal progenitors and brain endothelial cells (Bracko et al., 2012; Ferron et al., 2015; Logan et al., 1994). However, there is considerable controversy in the field over which receptor mediates IGF-II actions in the adult CNS. Two reports concluded that IGF-II induces neural stem/progenitor cell proliferation through the IGF-1R either in primary neurosphere cultures (Ferron et al., 2015; Lehtinen et al., 2011) or in transit amplifying progenitors (Ferron et al., 2015; Lehtinen et al., 2011). Yet another report concluded that IGF-II promotes hippocampal learning and memory consolidation indirectly through its binding to the cation-independent mannose-6-phosphate (M6P)/IGF2R, generally considered a scavenger receptor for IGF-II and M6P tagged proteins but with no inherent signaling capability (Chen et al., 2011). While our *in vitro* studies have supported the hypothesis that IGF-II promotes NSC self-renewal through the IR-A (Ziegler et al., 2014; Ziegler et al., 2012), no studies have directly tested the function of the IR on NSC self-renewal *in vivo*. In the studies reported here, we used both *in vitro* and *in vivo* genetic approaches to delete the IR in NSCs to test its function in NSC self-renewal as well as to evaluate the downstream consequences on neurogenesis and behavior.

The IR also is important in tumor biology. The IR is over-expressed in thyroid tumors in particular, where it stimulates the self-renewal of cancer stem cells. Consistent with these findings, the IR, and specifically IR-A, is associated with a more aggressive, undifferentiated tumor phenotype in several types of solid tumors. Glioblastoma multiforme (GBM) is the most common type of malignant brain tumor in adults, accounting for 78% of all CNS tumors. Among the GBMs the proneural and mesenchymal subtypes are the hardest to treat with high incidences of relapse. A prevailing view is that these relapses are due to the presence of self-renewing cancer stem cells that are resistant to chemo and radiation therapies (Chandran et al., 2015).

Gong et al (Gong et al., 2016) showed that the IR is commonly expressed in surgical specimens of GBM patients and contributes to GBM cell survival and growth through Akt dependent signaling. Moreover, an ATP-competitive IGF1R/IR inhibitor has shown promising results in decreasing cell viability and migration both *in vitro* and *in vivo* in temozolomide-resistant gliomas (Zhou, 2015). Based on our previous studies showing that IGF-II signaling through the IR promotes the self-renewal of NSCs in vitro and on data from other investigators showing that an IGF-II/IR signaling loop enriches for cancer stem cells in other tumors (Sciacca et al., 1999; Vella et al., 2002), we tested the hypothesis that IR and IGF-related genes are expressed at high levels in the most aggressive GBM subtypes and regulate growth of GBM cancer stem cells.

## Results

### IR deletion decreases SVZ-derived NSCs and increases progenitor lineages

IR deletion in neural stem/progenitor cells (NSPs) in vitro through adenovirus-Cre expression resulted in a 74% decrease in secondary neurosphere number (p=0.0001, Supplemental Fig. 1F) and a 78% decrease in neurosphere size between NSPs infected with Ad-Cre-GFP compared to control Ad-GFP virus (p<0.0001, Supplemental Fig. 1G). A measurement of transfection efficiency at 24 hrs post-infection revealed no significant differences between the Ad-Cre-GFP and control Ad-GFP infected populations (Supplemental Fig.1E). To determine whether deleting the IR reduced the frequency of NSCs, we established a ΔNSC-IRKO mouse line. Nestin-CreERT2 mice, where Cre expression is driven by the 2^nd^ intron of the nestin promoter (https://www.jax.org/strain/016261), were crossed with IR^loxP^ mice, where exon 4 of the IR is flanked by loxP sites (B6.129S4(FVB)-Insr^tm1Khn^/J) and that had previously been mated with a TdTomato reporter mouse line (B6.Cg-Gt(ROSA)26Sortm14(CAG-tdTomato)Hze/J), to generate NesCreERT2 ^+/-^ Tdt ^+/+^ IR ^fl/fl^ mice. As the 2^nd^ intron of the nestin promoter is active only in NSCs, only the NSCs will have the deletion of exon 4 causing a frame shift mutation leading to the immediate stop of translation. NestinCreERT2 ^-/+^ TDT ^+/+^ IR ^fl/fl^ (hereto referred to as ΔNSC-IRKO) and NestinCreERT2 ^-/+^ TDT ^+/+^ IR ^WT^ (ΔNSC-IR WT; controls) SVZ cells from P4 pups were grown *in vitro* as neurospheres and treated with 4-OH tamoxifen (0.5 µM) or vehicle for 24 hours to induce Cre transcription. The treated cells were propagated for an additional 5 days and then dissociated for flow cytometric analysis as outlined in Fig. 1A. Vehicle treated cells were tdTomato negative whereas tamoxifen treated cells were strongly tdTomato positive (Fig. 1B,B’). Using a flow cytometry panel established previously in our lab, cells within the neurosphere were parsed into NSCs and 7 different intermediate progenitors (Fig. 1C). This analysis revealed that the ΔNSC-IRKO spheres had 3.5-fold fewer NSCs compared to the ΔNSC-IR WT where only the Tdtomato+ cells were evaluated.

**Figure 1:**
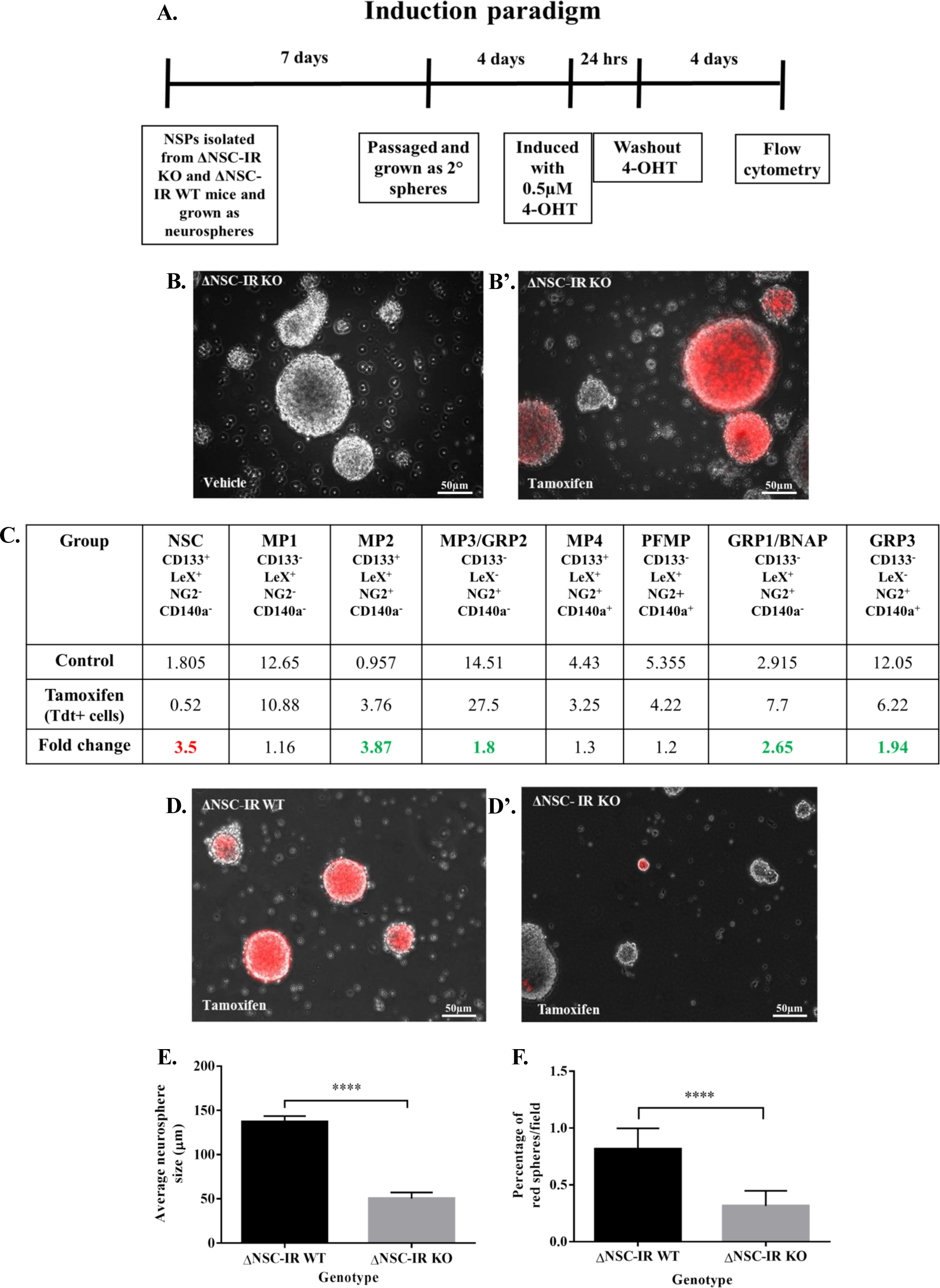
IR deletion in NSCs in vitro decreases NSCs and alters lineage. Induction paradigm of experiment. Cells from ΔNSC-IR WT and ΔNSC-IR KO mice were treated with 0.5 µM 4-OH tamoxifen or vehicle for 24 h. **B,B’.** Neurospheres from vehicle and tamoxifen treated cells 5 days after induction. **C.** ΔNSC-IR KO mice have fewer NSCs and a greater proportion of intermediate progenitors within the neurospheres than WT mice (red numbers indicate decreases in fold change, green numbers indicate increases in fold change). **D,D’.** Representative images of tertiary spheres after plating to measure sphere forming ability. **E.** Average sizes of tertiary Tdtomato red+ neurospheres. **F.** Percentage of tertiary Tdtomato red+ neurospheres per field were reduced in ΔNSC-IR KO mice (unpaired t test; p<0.0001). Data are representative of two independent experiments.

Correspondingly, the ΔNSC-IRKO had increased progenitor populations: MP2 (3.8-fold), MP3 (1.8-fold), GRP1 (2.6-fold) and GRP3 (1.9-fold) (Fig. 1C). These changes were not due to altered expression levels for the cell surface markers as levels of CD133, LeX, NG2 and CD140a were not different between IR WT cells; vehicle treated cells (tdT negative), ΔNSC-IR WT (tdT positive and negative cells) and ΔNSC-IRKO tdT negative cells (Supplemental Fig. 2). A subset of the cells used for the flow cytometric analysis was re-plated to analyze sphere-forming ability. Consistent with an essential role for IR in NSC self-renewal, fewer Tdtomato+ spheres formed in the ΔNSC-IRKO compared to ΔNSC-IR WT (Fig. 1 D-F).

**Figure 2:**
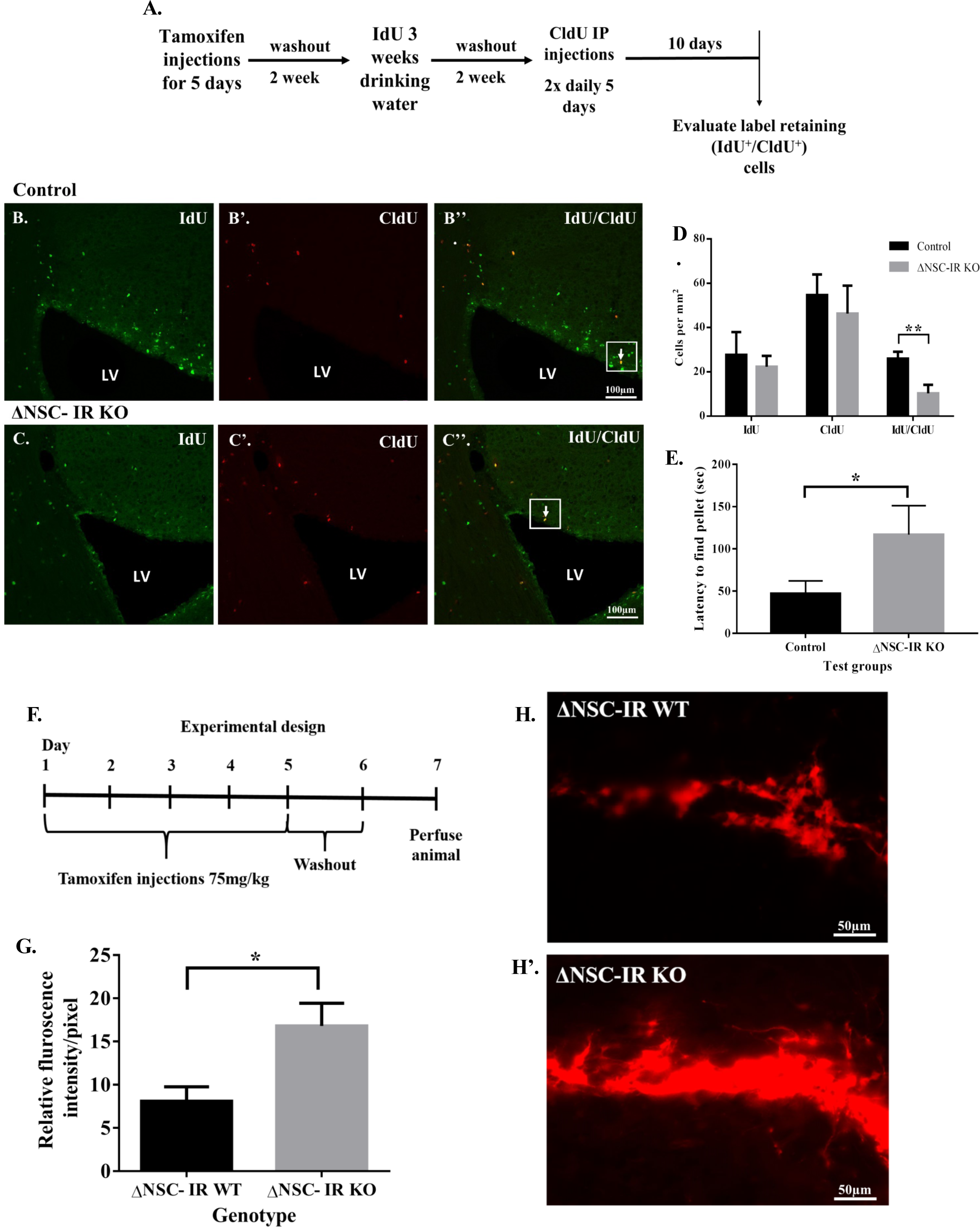
ΔNSC-IR KO mice have decreased numbers of SVZ NSCs, increased numbers of TdTomato+ SVZ progenitors and impaired olfaction. Schematic of experimental timeline depicting timing of tamoxifen and IdU and CldU administration. **B-B’’.** Representative images of IdU and CldU labeling in control SVZ. **C-C’’.** Representative images of IdU and CldU labeling in ΔNSC-IRKO SVZ. White arrows represent IdU^+^/CldU^+^ double positive cells (scale bar represents 100 µm). **D.** Numbers of IdU, CldU single and double positive cells that were located within 80 µm of the lateral ventricle were reduced by 50% in the ΔNSC-IRKO SVZ (Mean ± SEM, unpaired t test *p=0.03, n=4 control, n=5 IRKO). **E.** ΔNSC-IR KO mice required a significantly greater amount of time to find a buried food pellet compared to WT mice (Mean ± SEM, Mann Whitney test *p=0.01, n=10 control, n=12 IRKO). **F.** Schematic representation of IR deletion paradigm. **G.** Relative SVZ fluorescence intensity in ΔNSC-IRKO and ΔNSC-IR WT mice (Mean ± SEM, * p <0.05, unpaired t test, n = 3 per group). **H,H’.** Representative images of Tdtomato expression in ΔNSC-IRKO and ΔNSC-IR WT mice.

### IR deletion decreases forebrain SVZ NSCs in vivo and compromises olfaction

To establish whether the IR is essential for NSC self-renewal *in vivo,* ΔNSC-IRKO and ΔNSC-IR WT mice were administered tamoxifen or vehicle and analyzed 2 days later (Supplemental Fig. 3A). TdT was strongly expressed by nestin+ cells of the SVZ, SGZ and in the alpha tanycytes of the hypothalamus in mice that were Cre+ (Supplemental Fig. 3C,C’-3F,F’) whereas tdT+ cells were not evident in the tamoxifen administered Cre- mice (Supplemental Fig.3B,B’).

**Figure 3:**
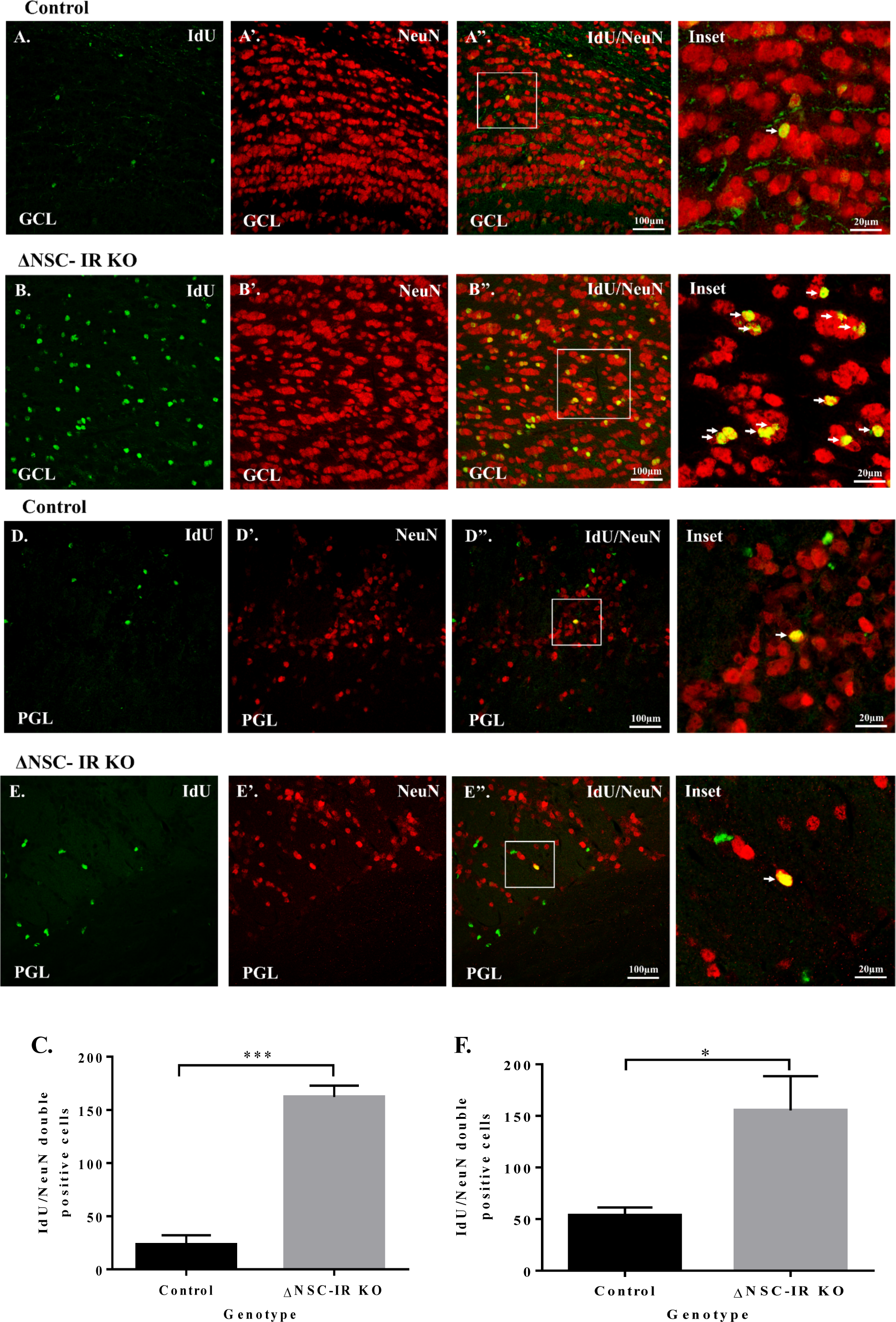
ΔNSC-IR KO mice have increased neurogenesis in the olfactory bulb. **A,A’,A’’.** Granule cell layer of control mice stained for IdU^+^(green)/NeuN^+^ (red) cells in the olfactory bulb 9 weeks after tamoxifen administration. **B,B’,B’’.** ΔNSC-IR KO mice stained for IdU^+^/NeuN^+^ (arrows) in the granule cell layer at 9 weeks after tamoxifen administration. **C.** Quantification of IdU/NeuN positive cells in the granule cell layer of control and ΔNSC-IRKO mice (Mean ± SEM, p < 0.001). **D,D’,D’’.** Periglomerular layer of control mice stained for IdU^+^(green)/NeuN^+^ (red) cells of the olfactory bulb 9 weeks after tamoxifen administration. **E,E’,E’’.** ΔNSC-IRKO mice showing increased IdU^+^/NeuN^+^ (arrows) in the periglomerular layer 9 weeks after tamoxifen administration. **F.** Quantification of IdU/NeuN positive cells in the periglomerular layer of control and ΔNSC-IRKO mice (Mean ± SEM, p < 0.05). Scale bars represent 100 µm in panels A-D and 20 µm for the insets. Images are representative of 3 animals per genotype.

To determine whether there were fewer label-retaining cells in the ΔNSC-IRKO neurogenic regions, iododeoxyuridine (IdU) and chlorodeoxyuridine (CldU) were administered at equimolar concentrations on different schedules over the course of 5 weeks (Fig. 2A) (Vega and Peterson, 2005). Using stereology methods, we counted the number of IdU and CldU single and double positive cells in the SVZ in ΔNSC-IRKO and ΔNSC-IR WT mice. This analysis revealed a 66% reduction in the number of IdU^+^/CldU^+^ double positive cells (NSCs) in the SVZ (cell numbers: ΔNSC-IR WT 26.5 ± 2.2 vs ΔNSC-IRKO 8.8 ± 2.3; unpaired t test p=0.03) (Fig. 2B-D). As neuroblasts produced within the SVZ perpetually regenerate interneurons in the olfactory bulbs we tested the mice for deficits in olfaction. Consistent with aberrant neurogenesis in the olfactory bulb, the ΔNSC-IRKO knockout mice took twice as long to find a hidden food pellet compared to the ΔNSC-IR WT mice (ΔNSC-IR WT 19.25 ± 3.1 seconds vs ΔNSC-IRKO 55.8 ± 19.2 seconds; Mann-Whitney test p=0.01), indicating deficits in olfactory sensitivity (Fig. 2E).

### Reduction in NSC number is not due to cell death

To assess whether the reduction in SVZ NSC number in the ΔNSC-IRKO was due to cell death, ΔNSC-IR KO and ΔNSC-IR WT mice were induced with tamoxifen for 5 days and sacrificed on day 6 (Fig. 2F). Cell death in the ΔNSC-IRKO would result in a reduction in the numbers of TdTomato+ cells. However, we observed no reduction in tdT+ cells. Rather, the density of tdT+ cells increased in the ΔNSC-IRKO compared to the ΔNSC-IR WT mice (ΔNSC- IR WT 8.1 ± 1.7 fluorescence intensity/pixel vs ΔNSC-IRKO 17 ± 2.6; unpaired t test p<0.05) (Fig. 2G-H’) (p = 0.018).

### IR deletion in NSCs increases proliferation of SVZ NSCs and production of olfactory bulb granule cell and periglomerular neurons

As described above, IR deletion compromised NSC self-renewal with a corresponding increase in several progenitor cell populations and a net increase in tdT+ cells within the SVZ. Consistent with the increase in progenitors, there was an increase in the numbers of IdU^+^/NeuN^+^ double positive cells in the granule cell layer (ΔNSC-IR WT 23.3 ± 8.7 cells vs ΔNSC-IRKO 162.2 ± 10.4; unpaired t test p<0.001) (Fig. 3 A-A”, B-B”, C) and in the periglomerular layer (ΔNSC-IR WT 53.6 ± 7.6 cells vs ΔNSC-IRKO 155.5 ± 33; unpaired t test p<0.05) (Fig. 3 D-D”, E-E”, F) of the olfactory bulb (OB) in the ΔNSC-IRKO vs ΔNSC-WT mice when analyzed 9 weeks after IR deletion.

### Hippocampal neurogenesis and hippocampal-dependent behaviors are unchanged in ΔNSC-IRKO mice

To establish whether IR is required to maintain the NSCs of the hippocampal SGZ, stereology was performed to count IdU single, CldU single and IdU^+^/CldU^+^ double labeled cells within the SGZ. In contrast to the SVZ, similar numbers of labeled cells were observed in both ΔNSC-IRKO vs ΔNSC-WT mice (Supplemental Fig. 4 A-C). To assess hippocampal spatial learning, the mice were evaluated using the Morris water maze test. Neither the acquisition rate, nor the total time spent in the quadrant that contained the platform during the probe test was different between the ΔNSC-IRKO vs ΔNSC-IR WT mice (ΔNSC-IR WT 24.5 ± 1.8 seconds vs ΔNSC-IRKO 27 ± 4.1) (Supplemental Fig. 4D). ΔNSC-IRKO mice were further evaluated for performance in the elevated plus maze to determine whether function of the ventral hippocampus was altered with deletion of the IR in the NSCs. There was no difference in the amount of time that ΔNSC-IRKO and ΔNSC-IR WT mice spent in the open arms of the maze (ΔNSC-IR WT 60.4 ± 10.1 seconds vs ΔNSC-IRKO 46.7 ± 12.7) (Supplemental Fig. 4E).

**Figure 4:**
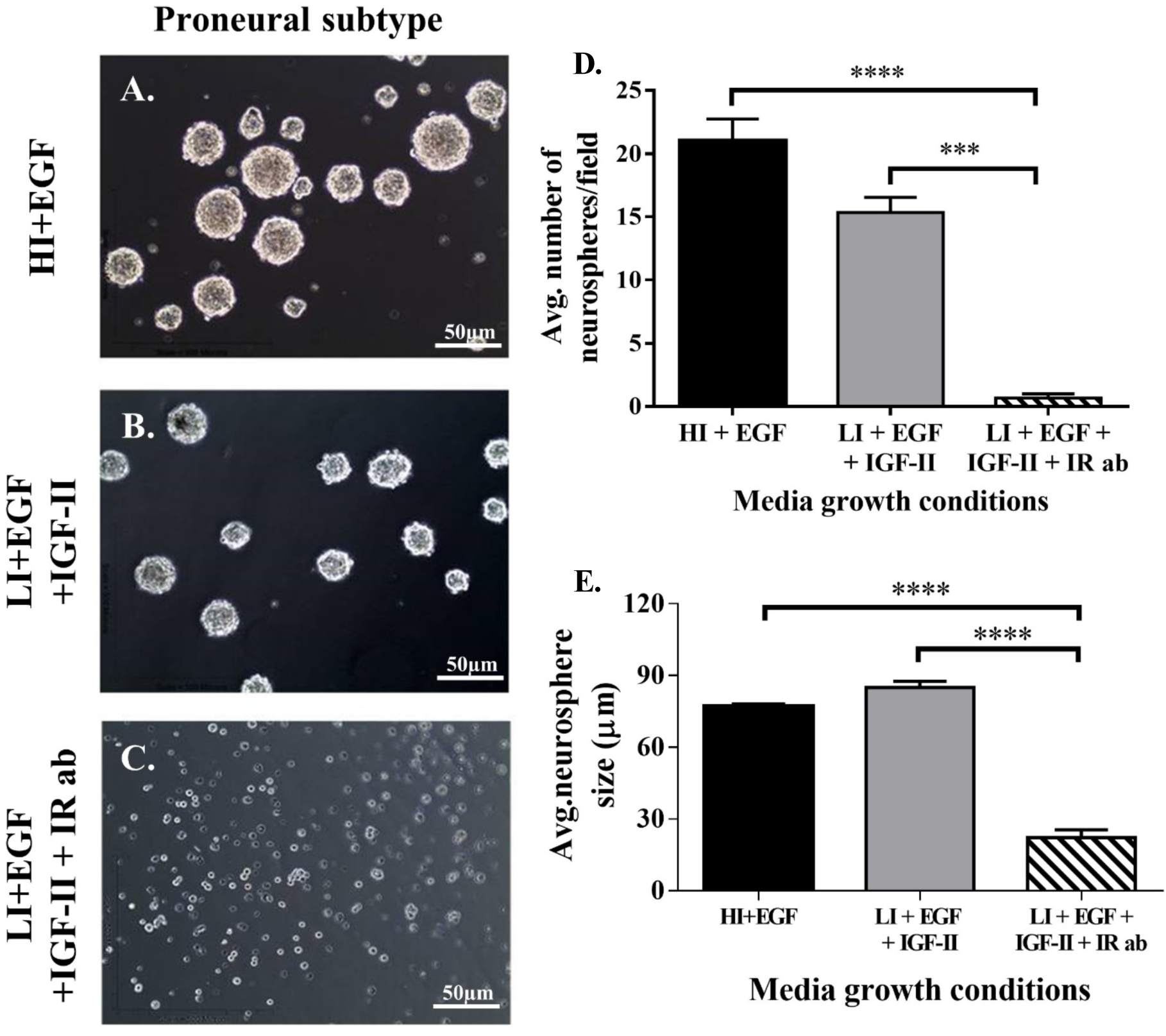
A function blocking antibody to the IR decreases self-renewal and proliferation of proneural GBM tumorspheres. **A.** Representative images of proneural GBM tumorspheres grown in defined culture medium. **B.** Representative images of proneural GBM tumorspheres grown in defined medium with physiological concentrations of insulin but supplemented with IGF-II (LI+EGF+IGF-II). **C.** Representative images of proneural GBM tumorspheres grown in LI+EGF+IGF-II with a function blocking IR antibody. **D.** Average number of tumorspheres/field ± SEM after 7 days of growth under different conditions (***p<0.001, ****p<0.0001). **E.** Average sizes of tumorspheres/field ± SEM, after 7 days of growth under different conditions (***p<0.001, ****p<0.0001). Data are representative of 3 independent experiments. Scale bar represents 50 µm.

Label retaining α-tanycytes have been reported along the walls of the 3^rd^ ventricle that produce new neurons and glial cells in the paraventricular, supraoptic and arcuate nuclei of the hypothalamus (Robins et al., 2013). Within the median eminence of the hypothalamus there is another set of progenitors that can generate neurons involved in metabolism (Lee et al., 2012). Therefore, we analyzed the relative numbers of IdU, CldU and double labeled cells in the hypothalamus and median eminence of the ΔNSC-IRKO vs ΔNSC-IR WT mice. IdU^+^ and CldU^+^ cells were much sparser in the hypothalamus than in the SVZ and the SGZ and stereological counts of cells in the lateral wall of the 3^rd^ ventricle failed to reveal any change in the number of α-tanycytes between the ΔNSC-IRKO vs ΔNSC-IR WT mice (Supplemental Fig. 5A-C). As the α-tanycytes can give rise to the neurons in the arcuate nucleus which controls food intake, we measured the body weight over the 5-day course of tamoxifen injection. Although there was a trend towards decreased body weight in ΔNSC-IRKO vs control mice, the difference did not reach statistical significance (Supplemental Fig. 5D). In the median eminence both the β2- tanycytes and rapidly proliferating CldU^+^ transit-amplifying cells showed a trend towards a decrease in number (Supplemental Fig. 6A-C).

**Figure 5:**
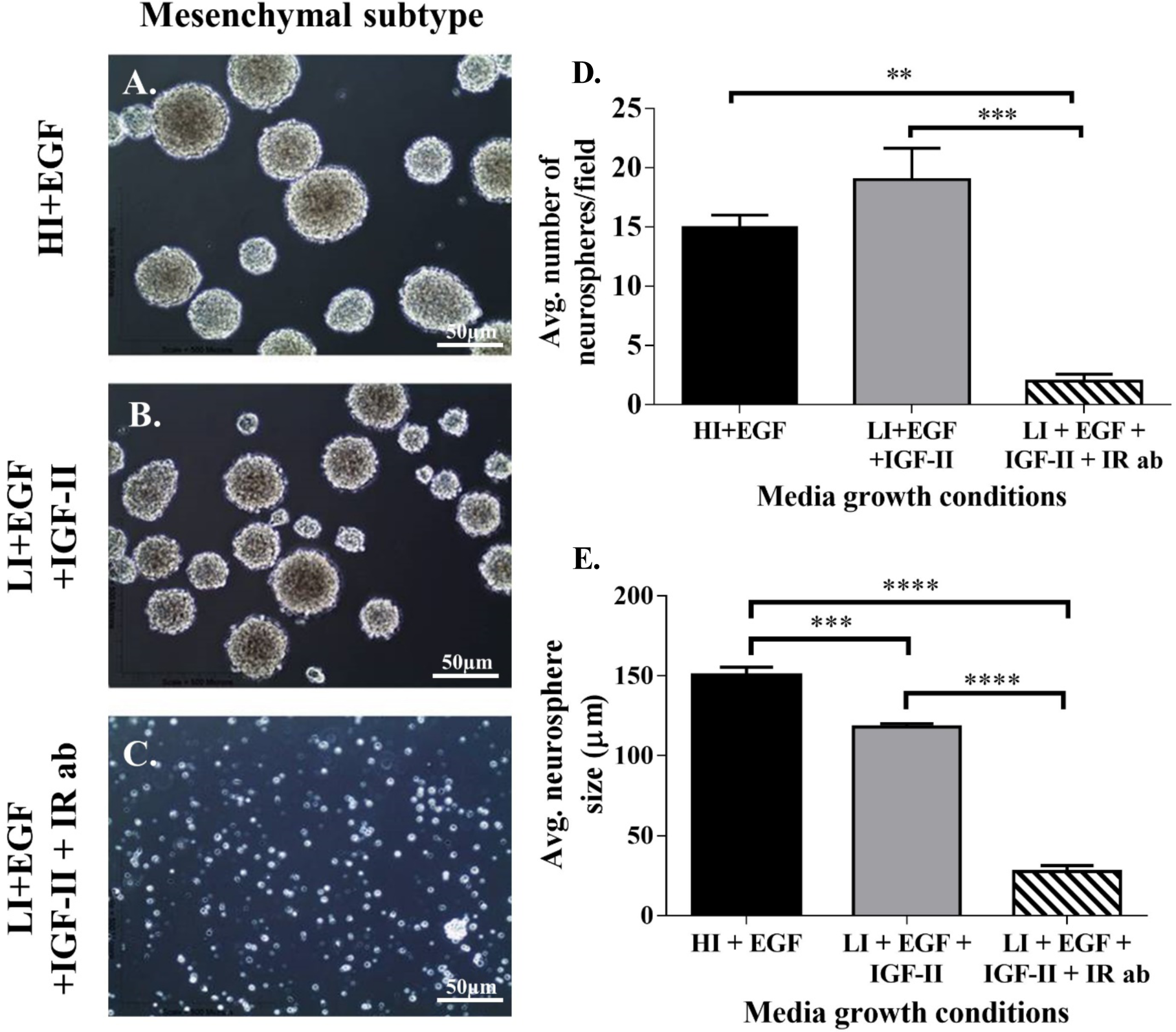
A function blocking antibody to the IR decreases self-renewal and proliferation of mesenchymal GBM tumorspheres. **A.** Representative images of mesenchymal GBM tumorspheres in medium with superphysiological levels of insulin (HI+EGF). **B.** Representative images of mesenchymal GBM tumorspheres grown with physiological levels of insulin+IGF-II (LI+EGF+IGF-II). **C.** Representative images of mesenchymal GBM tumorspheres grown in LI+EGF+IGF-II with a function blocking IR antibody **D.** Average number of tumorspheres/field ± SEM after 7 days of growth under different conditions (**p<0.01, ***p<0.001, ****p<0.0001). **E.** Average size of tumorspheres/field ± SEM after 7 days of growth under different conditions (***p<0.001, ****p<0.0001). Data are representative of 3 independent experiments. Scale bar represents 50 µm.

### In silico data analysis reveals elevated expression of insulin/IGF genes and splice enzymes in proneural and mesenchymal subtypes of GBM

A number of studies have implicated overexpression of IR/IGF related genes in cancer (Chettouh et al., 2013; Kim et al., 2012; Vella et al., 2001; Vigneri et al., 2015; Wang et al., 2013). Therefore, we queried expression data for genes in the IR/IGF growth factor system from the TCGA database for proneural (p) (n=31) and mesenchymal (m) (n=55) GBM subtypes. The proneural and mesenchymal GBM subtypes had higher expression of the IR; proneural (5.6 fold), mesenchymal (4.9 fold), IGF1R; proneural (10.4 fold), mesenchymal (11.3 fold), and IGF2; proneural (59 fold), mesenchymal (60 fold) when compared to normal astrocytes (Table 1). Chettouh et al. 2013 showed that in hepatocellular carcinoma the change in ratio of IR-A to IR-B was due to alterations in splice enzymes. Therefore, we queried the expression of the enzymes known to be involved in splicing the mRNA for the IR. In agreement with the earlier paper, we observed increased expression of several splice enzymes involved in IR alternative splicing. (CUCGBP1; proneural (4.9 fold), mesenchymal (4.5 fold), HNRNPHI; proneural (2.9 fold) mesenchymal (2.3 fold), HNRNPA2B1; proneural (2.5 fold), mesenchymal (2.1 fold), SFRS1/AS2; proneural (2.8 fold), mesenchymal (2.4 fold) (Table 1).

**Table 1:** Insulin/IGF growth factor system genes and IR mRNA splicing factor genes are induced in Glioblastomas. 1,2,3. TCGA data mining for genes involved in the insulin/IGF growth factor system reveal higher expression of IR, IGF-1R and IGF2 in proneural and mesenchymal subtypes vs. normal astrocytes (proneural n=31, mesenchymal n=55). **3,4,5,6.** TCGA data mining for enzymes involved in IR mRNA splicing in proneural and mesenchymal subtype (proneural n=31, mesenchymal n=55). Data represents mean of all values.

| No. | Gene name | Fold change in Expression Proneural | Fold change in Expression Mesenchymal |
| --- | --- | --- | --- |
| 1 | INSR | 5.6 | 4.9 |
| 2 | IGF-1R | 10.4 | 11.3 |
| 3 | IGF2 | 59 | 60 |
| 4 | (CUCGBP1)<br>CELF1 | 4.9 | 4.5 |
| 5 | HNRNPH1 | 2.9 | 2.3 |
| 6 | HNRNPA2B1 | 2.5 | 2.1 |
| 7 | SFRS1 | 2.8 | 2.4 |

### Proneural and mesenchymal subtype glioblastoma stem cells switch IR-A/IGF1R ratio

Our prior studies demonstrated that 1) IGF-II promotes stemness in adult neural stem cells (Ziegler et al., 2012)(Ziegler 2019), 2) IGF-II mediates SVZ NSC self-renewal through IR in vitro (Ziegler et al., 2014), and 3) the IR-A is highly expressed in the medial SVZ (Ziegler et al., 2014). Thus, we performed studies to establish whether gene expression changes seen in entire tumors reflected changes occurring within the GBM stem cells. Multiple studies in different cancer types have shown a switch in the ratio of IR-A/IR-B and IR-A/IGF-1R that allow cancer cells to shift from a metabolic to a more proliferative phenotype (Andres et al., 2013; Chettouh et al., 2013; Garofalo et al., 2013; Lodhia et al., 2015; Ulanet et al., 2010). We generated tumorspheres from proneural and mesenchymal subtype human tumors and performed an RNA transcriptome analysis. Again, the ratio of the IR to the IGF-1R was 4.9-fold higher in the proneural GBM spheres and 3-fold higher in mesenchymal GBM spheres compared to normal human neural stem cells (Table 2). Interestingly, IR-B (which contains exon 11) was not detected in either normal neurospheres or tumorspheres consistent with our prior findings in the mouse SVZ *in vivo* (Ziegler et al., 2014).

**Table 2:** IR-A/IGF-1R ratio Increases in Proneural and Mesenchymal GBM stem cells. . Insulin receptor (INSR) and IGF-1R mRNA levels were quantified from GBM neurospheres from deep sequencing using Roche 454/GS FLX Sequencing Technology. Exon 11 was not detected in the NSC populations (GB2, WCR8, HNSC). Thus, the INSR in NSCs corresponds to IR-A (Italics indicates increased expression). (Proneural n=2, mesenchymal n=2, normal human neural stem cells n=2). Data represents mean of values.

| **Gene** | **Avg. of hNSC**<br>**spheres** | **Avg. of**<br>**Proneural**<br>**Spheres (GB2)** | **Avg. of**<br>**Mesenchymal**<br>**spheres**<br>**(WCR8)** |
| --- | --- | --- | --- |
| **IGF-1R** | **41.5** | **6.48** | **11.49** |
| **INSR** | **7.29** | **5.44** | **6.06** |
| **Ratio of**<br>**INSR/IGF-1R** | **0.17** | **0.83** | **0.52** |
| **Fold change** | **1** | **4.9** | **3** |

### Blocking the IR alters the number and size of proneural and mesenchymal GBM tumorspheres

Sphere-forming assays have been widely used to evaluate the self-renewal properties of stem cells, including cancer stem cells, using repeated passage as an index of self-renewal. Our lab previously showed that lowering the concentration of insulin to physiological levels (4.4 nM) and adding IGF-II (224 ng/ml), increased numbers of murine SVZ neurospheres formed upon subsequent passages, consistent with enhanced stemness. Therefore, we asked whether lowering the concentration of insulin and adding IGF-II to the medium used to propagate human tumorspheres enhance GBM tumorsphere growth. Contrary to our findings with normal neurospheres grown in vitro, there was no difference in the number of proneural or mesenchymal GBM tumorspheres formed when the cells were grown in low insulin medium supplemented with IGF-II compared to control conditions containing high insulin likely due to the high endogenous expression of IGF-II in the GBMs. However, addition of an IR blocking antibody resulted in a 35-fold reduction in the number of proneural tumorspheres formed in EGF supplemented medium (HI+EGF 21 ± 1.7 vs LI+IGF-II+ IR ab 0.6 ± 0.3; One-way ANOVA p<0.0001) and an 18.5 fold reduction in the number of tumorspheres in EGF + IGF-II supplemented medium (LI+ EGF + IGF-II 11.1 ± 1.2 vs LI+ EGF +IGF-II+ IR ab 0.6 ± 0.3; One-way ANOVA p<0.001) (Fig. 4 A-D). In mesenchymal tumorspheres supplemented with the IR blocking antibody we observed a 7.5 fold reduction in tumorsphere number compared to media with EGF (HI+EGF 15 ± 1 vs LI+IGF-II+ IR ab 2 ± 0.5; One-way ANOVA p<0.01) and a 9.5 fold reduction in tumorsphere number compared to media supplemented with EGF + IGF-II (LI+ EGF + IGF-II 19 ± 2.6 vs LI+ EGF +IGF-II+ IR ab 2 ± 0.5; One-way ANOVA p<0.001) formed upon passage (Fig. 5 A-D). These data support the conclusion that the IR is essential to maintain the stemness of proneural and mesenchymal GBM cancer stem cells.

In breast and colorectal cancers, IGF-1R promotes the proliferation of the progenitors within the tumor (Farabaugh et al., 2015; Rota et al., 2014). As shown in Table 2, there is a switch in the ratio of IGF-1R to IR-A in the GBM spheres compared to normal human neural stem cells. To establish whether the IR was promoting growth of the tumorspheres, proneural GBM tumorspheres were grown in medium containing the IR blocking antibody. When the sphere sizes were measured as an index of growth, inhibiting the IR reduced sphere sizes by 3.5 fold in EGF supplemented medium (HI+EGF 77 ± 1.1 vs LI+IGF-II+ IR ab 22 ± 3.17; One-way ANOVA p<0.0001) and by 3.8 fold in EGF + IGF-II media supplemented with the IR blocking antibody for the proneural subtype (LI+ EGF + IGF-II 85 ± 2.64 vs LI+ EGF +IGF-II+ IR ab 22 ± 3.17; One-way ANOVA p<0.001) (Fig. 4 A-C, E). Similarly, inhibiting the IR in the mesenchymal subtype reduced tumorsphere growth by 5.3 fold in EGF supplemented medium (HI+EGF 150 ± 5.5 vs LI+ EGF+ IGF-II+ IR ab 28 ± 3.7; One-way ANOVA p<0.0001) and by 4.2 fold in EGF + IGF-II media supplemented with the IR blocking antibody (LI+ EGF+ IGF-II 118 ± 2.03 vs LI+ EGF +IGF-II+ IR ab 28 ± 3.7; One-way ANOVA p<0.0001) (Fig. 5 A-C, E). The addition of a non-specific IgG had no effect on tumorsphere size or number (Supplemental Fig. 7).

## Discussion

Although the IR-A isoform of the IR is known to regulate NSCs in vitro and is elevated in several tumor types, IR function in regulating NSC homeostasis in vivo has not been previously investigated. The findings reported here demonstrate that the IR is necessary to maintain a subset of normal adult NSCs. Deleting the IR in SVZ-derived NSPs in vitro decreased the numbers of NSCs with a corresponding increase in a subset of multipotential and bipotential neural progenitors. Consistent with the reduction in NSCs, IR deletion reduced the numbers of 2° neurospheres after passage. Deleting the IR *in vivo* similarly reduced the number of NSCs located in the medial SVZ with a corresponding expansion in neural progenitors accompanied by olfactory deficits. Interestingly, our findings revealed that the IR uniquely regulates NSCs in the SVZ since NSC numbers in the SGZ of the hippocampus and along the lateral wall of the 3^rd^ ventricle (α-tanycytes) were unaffected by loss of the IR. Altogether our results demonstrate that the IR is an essential receptor for self-renewal of a subset of NSCs and that its loss results in their depletion with functional consequences.

### IR-A is necessary to exert effects of IGF-II on adult stem cell self-renewal

Our prior data strongly support the conclusion that it is the IR-A isoform, and not the metabolic IR-B isoform, that is responsible for SVZ NSC self-renewal. Laser microdissection studies revealed that the IR-A is more highly expressed than the IGF-1R within medial aspect of the SVZ, whereas the IGF-1R is more highly expressed than IR-A in the lateral SVZ, which is the progenitor-cell domain of the SVZ (Ziegler et al., 2012). By contrast, the IR-B was undetectable in the SVZ. The IR-A also is the predominant isoform expressed in neurospheres and, as cells become lineage-restricted, the levels of IR-A decrease (Ziegler et al., 2012). There are numerous lines of evidence supporting the conclusion that IGF-II is the endogenous ligand that signals through the IR-A to maintain the NSCs in the adult SVZ. Our prior studies showed that IGF-II promotes stemness of SVZ-derived NSCs *in vitro* through the IR-A. Moreover, the alterations resulting from our *in vivo* deletion of the IR in SVZ NSCs largely phenocopies the effect of IGF-II deletion from the adult SVZ (Ziegler et al., 2019). IGF-II deletion decreased NSCs in the SVZ, expanded the transit amplifying progenitors and caused olfaction deficits. These data support the conclusion that an IGF-II/IR-A signaling loop regulates NSC homeostasis (Ziegler et al., 2014; Ziegler et al., 2019; Ziegler et al., 2015; Ziegler et al., 2012). Although insulin also binds to the IR-A with high affinity, the biological effects of insulin versus IGF-II on IR intracellular signaling have not been studied in NSCs. Studies using murine fibroblasts engineered to express only IR-A or IR-B have established that the gene expression profiles for the two ligands differ (Pandini et al., 2003). Other studies have shown that activating the IR-A stimulates cell proliferation and promotes survival; whereas the IR-B regulates metabolism (Belfiore et al., 2009; Belfiore and Malaguarnera, 2011; Sacco et al., 2009; Sciacca et al., 2003). Recently, the mechanism for IGF-II binding to IR-A was elucidated (Alvino et al., 2011; Andersen et al., 2017; Rajapaksha et al., 2012). These data point to a role for IGF-II via IR-A that is distinct from the actions of insulin (Rajapaksha and Forbes, 2015). Using the neurosphere assay to analyze NSC self-renewal across passage, we demonstrated that IGF-II, but not insulin or IGF-I, maintained the SVZ NSCs and also expanded their numbers suggesting that IGF-II promotes symmetrical vs. asymmetrical divisions (Ziegler et al., 2012).

### IGF-II acts through both the IGF-1R and the IR to support adult neural stem cell self-renewal

It is becoming increasingly clear that there is not a single NSC, but rather that multiple NSC subtypes exist in the adult CNS. The lack of a phenotype in the SGZ NSCs with IR deletion provides yet another example of NSC heterogeneity. Our prior data taken together with the data presented here show that IGF-II acts through the IR-A to maintain SVZ NSCs whereas the IR is dispensable for SGZ NSC self-renewal. There are a number of studies that show the IGF-II is important for hippocampal neurogenesis (Ferron et al., 2015; Ouchi et al., 2013; Tao et al., 2015). Our recent characterization of adult neurogenesis in the conditional IGF-II KO showed that the number of NSCs in the SGZ declined when IGF-II was deleted. Moreover, depleting IGF-II in adulthood also caused deficits in spatial learning and increased anxiety (Ziegler et al., 2019). Bracko et al. 2012 (Bracko et al., 2012), suggested that SGZ NSC self-renewal is regulated by IGF-II via IGF-1R and AKT-dependent signaling. Consistent with their findings, deleting the IR had no effect on label-retaining cells in the SGZ or on spatial learning and memory ability or anxiety. It should be noted that the original genetic experiments on IGF-II demonstrated that this growth factor promotes fetal growth through both the IGF-1R and IR (Baker et al., 1993; Liu et al., 1993; Louvi et al., 1997). Altogether these data support the conclusion that IGF-II acts through the IR in the SVZ whereas it signals through the IGF-1R in the SGZ.

### There are multiple types of adult stem cells in the SVZ

Accumulating evidence shows that there are multiple subtypes of NSCs within the adult SVZ. Here we show that deleting the IR within the NSPs *in vitro* decreased the NSC population by ∼70%. Similarly, IR deletion from the NSCs *in vivo* resulted in a ∼70% reduction in label-retaining cells in the SVZ. Interestingly, conditionally deleting IGF-II in the adult mouse produced a ∼50% decrease in the percentage of NSCs (Ziegler et al., 2019). Altogether these observations indicate that IGF-II signaling via the IR is necessary to maintain a subset of NSCs within the adult SVZ. Other studies have described subsets of NSCs within the SVZ. Azim et al. 2014, showed that glutamatergic neuronal progenitors and oligodendrocyte precursors are derived from the dorsal SVZ under the control of Wnt/β-catenin signaling, while GABAergic neural precursors are derived from the lateral/ventral SVZ and are independent of Wnt/β-catenin signaling (Azim et al., 2014). Merkle et al., 2017 demonstrated that spatially segregated NSPs give rise to different olfactory bulb neurons (Merkle et al., 2007). Using Ad-Cre-GFP virus to label SVZ radial glia dorsally and ventrally, they showed that the NSCs in the dorsal region produced tyrosine hydroxylase positive (TH+) periglomerular cells while the ventrally located labeled NSCs gave rise to Calbindin positive (CalB+) periglomerular neurons. These data support the conclusion that there is significant functional heterogeneity amongst the SVZ NSCs.

### IR-A signaling in SVZ NSCs is not necessary for cell survival

The IR is important for the survival of cells throughout the body; therefore, the loss of the IR in the NSCs might have resulted in their premature demise. However, instead of NSC loss, IR deletion led to a loss of stemness in the neurosphere assay with an increase in the more rapidly dividing progenitors. These findings were corroborated by flow cytometric experiments that revealed a decrease in NSC number with a compensatory increase in the other intermediate progenitors, indicating that decreased NSC number observed was not due to cell death. Similarly, in vivo deletion of the IR further demonstrated that SVZ NSCs were reduced with a corresponding increase in neurogenesis.

### The IGF-II/IR-A signaling loop regulates stem phenotypes in tumor CSCs

Accumulating evidence over the last few decades has shown that the IR is abnormally expressed in many malignancies such as breast, colon, lung and thyroid (Chettouh et al., 2013; Kim et al., 2012; Vella et al., 2001; Vigneri et al., 2015; Wang et al., 2013); the IR-A is often more highly expressed than IR-B in these cancers. A number of studies have reported an IGF-II/IR-A autocrine loop that drives tumor progression (Sciacca et al., 1999; Vella et al., 2002). IGF-II binding to IR-A does not cause internalization of the receptor unlike the binding of insulin, leading to prolonged mitogenic activation allowing cancer cell proliferation (Morcavallo et al., 2012). Other studies have shown that a switch in the ratio of IR isoforms leads to the malignant transformation of cancer cells. Chettouh et al. 2013 (Chettouh et al., 2013) showed that in hepatocellular carcinoma a switch in the ratio of IR-A/IR-B is caused by alterations in the splice enzymes involved in IR mRNA splicing. Our data mining results reveal that there is a similar overexpression of the enzymes involved in IR mRNA splicing in proneural and mesenchymal GBM, leading to higher expression of the mitogenic IR-A. This would contribute to an IGF-II/IR-A autocrine signaling loop to promote the self-renewal of GBM stem cells. In addition to documented changes in levels of these components of the IGF-II system in cancer, there also are a number of studies that have implicated the IGF-2 mRNA binding proteins. These RNA binding proteins increase the stability of IGF-2 mRNA, resulting in production of higher levels of IGF-II protein. Interestingly, IGF-2 binding protein-2 is overexpressed in glioblastomas, and increased expression of IGF-2 binding protein-2 correlates with a poor prognosis for proneural GBMs (Cao et al., 2018).

Our findings here taken together with our previously published data support the conclusion that the IR-A is essential for maintenance of a subset of NSCs, and that the aggressive subtypes of GBM; proneural and mesenchymal, use an IGF-II/IR-A signaling loop to enhance self-renewal of tumor stem cells. These findings raise the possibility that IGF-II and IR- A should be considered as targets for new therapeutics either to enhance normal NSC function or to inhibit stem cell populations in GBMs.

## Materials and Methods

### Neurosphere propagation and quantification

Neurospheres were generated by enzymatically dissociating the periventricular region of IR^fl/fl^ pups (P4-5) as described previously (Ziegler et al., 2019) (see supplementary methods).

Adenovirus carrying the Cre recombinase gene with a GFP reporter (Vector Biolabs Cat #1700) or adenovirus carrying only GFP reporter (Cat #1060) (control) were infected at an MOI of 1000 during passage into secondary spheres. For in vitro induction of recombination, NestinCreERT2 ^- /+^ TDT ^+/+^ IR ^fl/fl^ generated secondary spheres were allowed to grow for 4 DIV and then 0.5 µM of 4-hydroxy tamoxifen (Sigma-Aldrich) or vehicle (ethanol) was added to the media for 24 hrs.

### Animals

All experiments were performed in accordance with protocols approved by the institutional animal care and use committee of Rutgers-New Jersey Medical School and in accordance with the National Institute of Health Guide for the Care and Use of Laboratory Animals (NIH Publications No. 80-23) revised in 1996. Tamoxifen (Sigma T5648) was dissolved in a corn oil: ethanol (9:1) and given I.P, at 75 mg/kg for 5 days. Mouse strains included CreER driven off a 2nd Intron promoter (Jax mouse stock # 016261) floxed IR mice (Jax mouse stock # 006955) and stop-flox Tdtomato mice (Jax mouse stock # 007905). Both male and female, heterozygous Cre mice, were used for the experiments. Progeny from breedings consisted of homozygous Tdtomato and homozygous floxed IR littermates that were either heterozygous Cre or Cre negative (wild-type, WT).

### Flow cytometry

Spheres were dissociated and counted as previously described (Buono et al., 2012) (see supplemental methods). For surface marker analysis, cells were incubated in PGB for 25 min with antibodies against Lewis-X (1:20, LeX/CD15 FITC, MMA; BD Bioscience), CD133-APC (1:50,13A4; eBioscience), CD140a (1:400, APA5; BioLegend) and NG2 Chondroitin Sulfate Proteoglycan (1:50, AB5320; Millipore). Goat anti-rabbit IgGAlexa Fluor 700 (1:100; Invitrogen) was used for NG2. All sample data were collected on the BD LSR II (BD Biosciences Immunocytometry Systems). Post-acquisition analysis was performed using FlowJoX (Tree Star Inc, Ashland, OR).

### Behavioral Tests

The buried food test for mouse olfaction and Morris water maze (hidden platform) were conducted as for our previous study(Ziegler et al., 2019) Elevated plus maze to test anxiety was performed according to Walf et al 2007 (Walf and Frye, 2007) (see also Supplementary Methods).

### Stereology

IdU / CldU administration and detection was performed as previously described (Ziegler et al., 2019) (see supplementary methods). The Olympus BX51 microscope was used to measure immunofluorescence of mounted slides of FB SVZ, SGZ and HT SVZ. Stereoinvestigator software was used to quantify the number of IdU^+^, CldU^+^ and IdU^+/^CldU^+^ double positive cells.

### Statistical analyses

All statistical analyses were performed using Graph Pad Prism software. For analysis of two groups, the unpaired t-test was used. For greater than two groups, One-way ANOVA with Tukey’s post hoc was used. In all cases, p value <0.05 was considered to be statistically significant.

### Subtyping GBM samples and Insilico data mining

GBM tumorspheres were obtained from Dr. Nikolaos Tapinos (Brown University). They were subtyped using a combination of pathology reports and a PCR mutation panel from SA Biosciences (**qBiomarker Somatic Mutation PCR Array: Human Brain Cancers).**

RNAseqV2 data matrix was obtained from the level 3 data of all batches available for public access in the TCGA database. Then for each TCGA barcode, the RSEM normalized genes file was downloaded. All these files were consolidated onto one excel spreadsheet. Using the supplemental information in Bernan et al. 2013, the files from each TCGA barcode were classified to the subtype of GBM that they belonged to proneural and mesenchymal GBM subtypes. Genes measured were normalized to RNA seq data of normal human astrocytes expression data from a Standford database (http://web.stanford.edu/group/barres_lab/brain_rnaseq.html).

### Expression analysis of RNAseq data

RNA was obtained from stem cell culturess isolated from two glioblastoma tumors (code-named GB2, WCR8) and from one normal human neural stem cell culture (HNSC). Next-generation RNA-sequencing was performed in the Epigenomics Core Facility of the Albert Einstein College of Medicine, NY, using an Illumina HiSeq2500 machine (see Supplemental Methods).

### Tumorsphere propagation and quantification

Human glioblastoma stem cells were provided by Dr. Nikolaos Tapinos lab (Brown University, Rhode Island). GBM stem cells from the two subtypes of GBM namely GB2; proneural, WCR8; mesenchymal were grown as described above. Neurospheres were collected after 5-7 days, dissociated and plated at 1 x 10^5^ cells/ml in the same media with human recombinant IGF-II (224 ng/ml), with IGF-II and IgG isotype antibody (Santa Cruz), or with IGF-II and IR blocking antibody. Neurospheres were quantified as previously described (157) using ImageJ software.

## Acknowledgements

This work was supported by R21 NS076874 awarded to SWL and TLW.

## Supplemental Figures and Text

**Supplemental figure 1.**
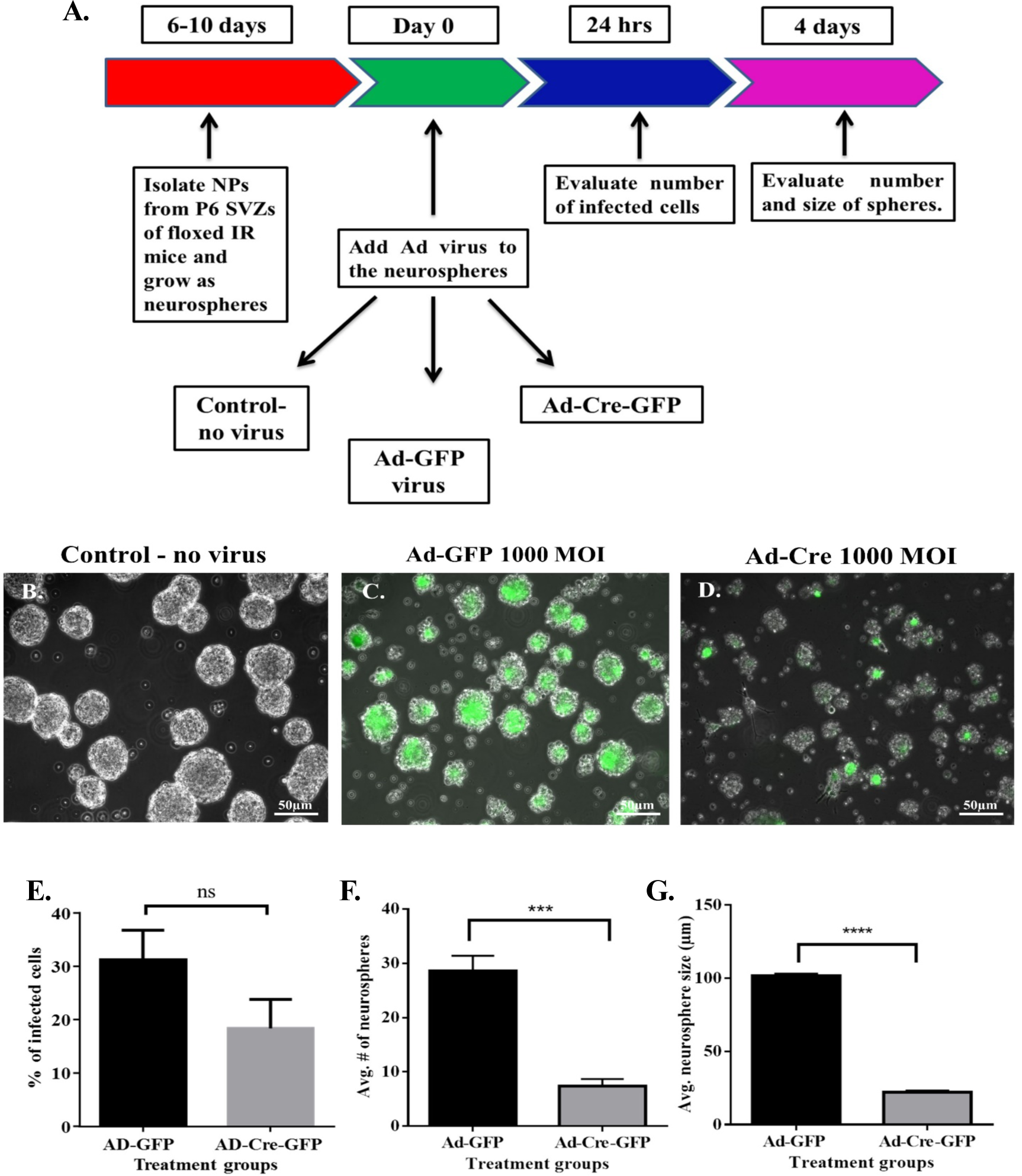
IR knockdown decreases neurosphere size and number of neural progenitors. Neurospheres generated from IR^fl/fl^ mice were dissociated into single cells and then infected with Ad-GFP, Ad-Cre or no-virus (control). Cells were grown under neurosphere-producing conditions. Images were captured 96hr after infection. **A.** Schematic of experimental paradigm. **B.** Control-no virus **C.** Spheres produced by cells infected with Ad-GFP virus **D.** Cells infected with Ad-Cre-GFP virus. Neurospheres infected with 1000 MOI Ad-GFP, Ad-Cre-GFP virus and control were counted and measured using the ImageJ software. **E.** Infection efficiency **F.** Average number of neurospheres. **G.** Average size of neurospheres. Mean ± SEM, ****p<0.0001,*** p=0.001 by unpaired t test. Scale bar represents 50 µm.

**Supplemental figure 2.**
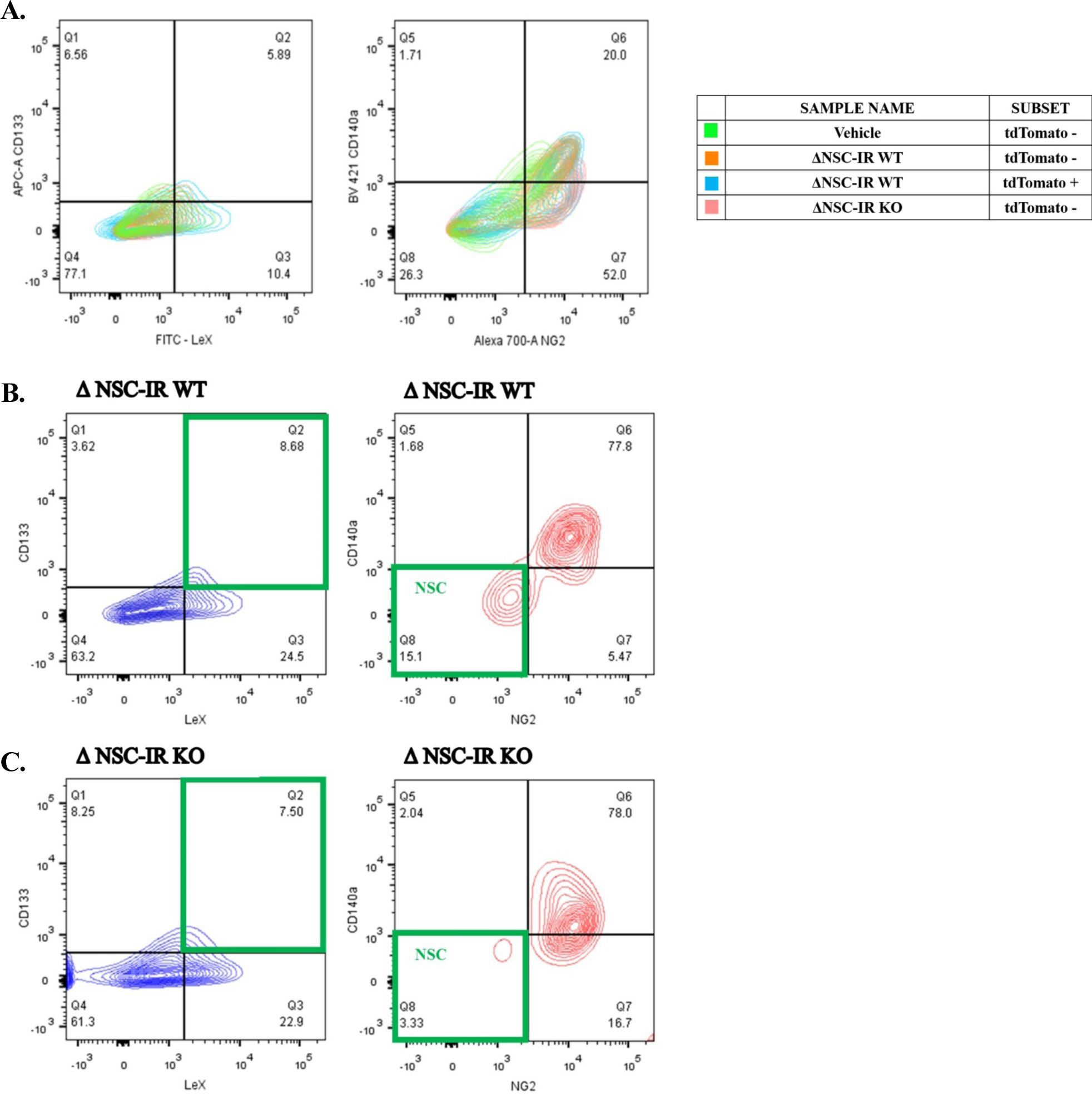
IR knockout decreases NSC population. Neurospheres generated from ΔNSC-IR WT and ΔNSC-IR KO mice were treated with 0.5 µM of 4-OH tamoxifen or vehicle for 24 hour followed by dissociation for flow cytometry. **A**. Control groups showing similar CD133, LeX, NG2 and CD140a expression. **B.** NSC numbers in ΔNSC-IR WT (control) induced with tamoxifen **C.** NSC numbers in ΔNSC-IR KO induced with tamoxifen. Representative data of one flow experiment.

**Supplemental figure 3.**
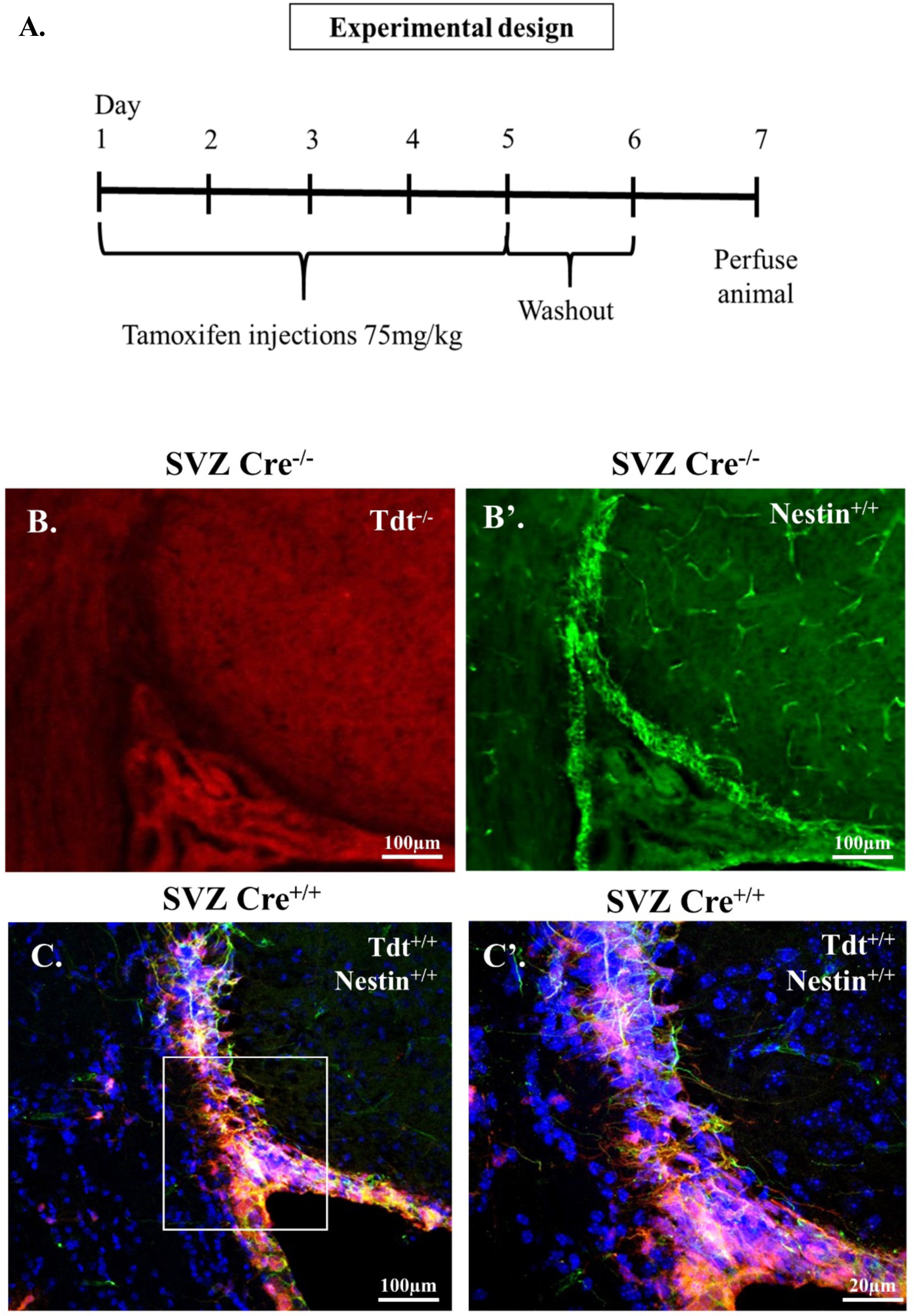
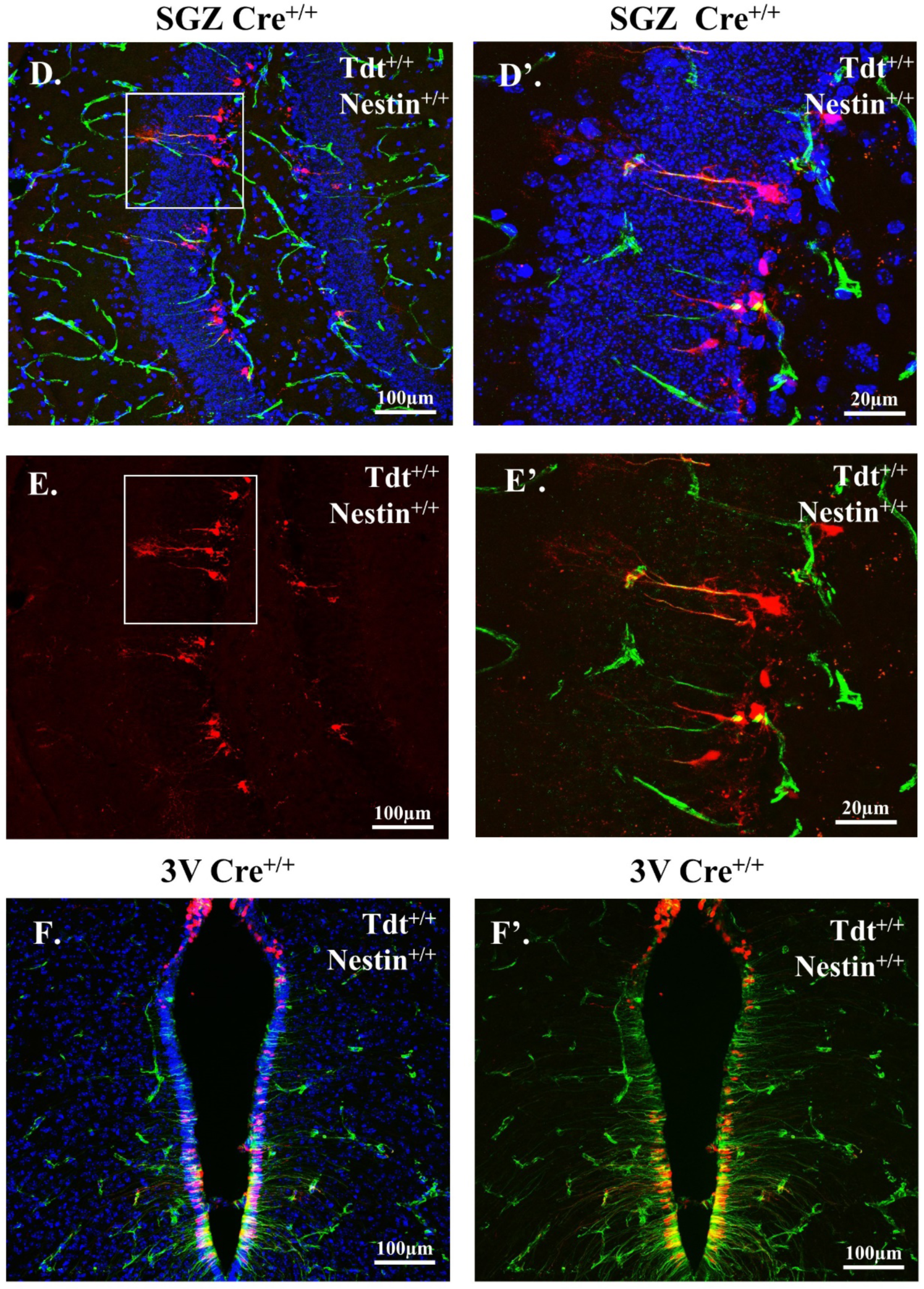
NestinCre promoter induces TdTomato expression in FB SVZ, SGZ and hypothalamus. **A.** Schematic of experimental timeline showing administration of tamoxifen and tissue collection. **B.B’.** SVZ of Cre negative mice induced with tamoxifen that are negative for tdTomato expression but stain positive for nestin with nestin antibody. **C.C’.** SVZ of Cre positive mice induced with tamoxifen showing tdTomato expression that colocalizes with nestin positive cells (yellow cells). **D.D’.** SGZ of Cre positive mice induced with tamoxifen showing tdTomato expression that colocalizes with nestin positive cells (yellow cells). **E.E’.** SGZ of Cre positive mice induced with tamoxifen showing tdTomato expression that colocalizes with nestin positive cells (yellow cells) without DAPI. **F.F’.** 3^rd^ ventricle of Cre positive mice induced with tamoxifen showing tdTomato expression that colocalizes with nestin positive cells (yellow cells) with and without DAPI. Scale bar represents 100 µm.

**Supplemental figure 4.**
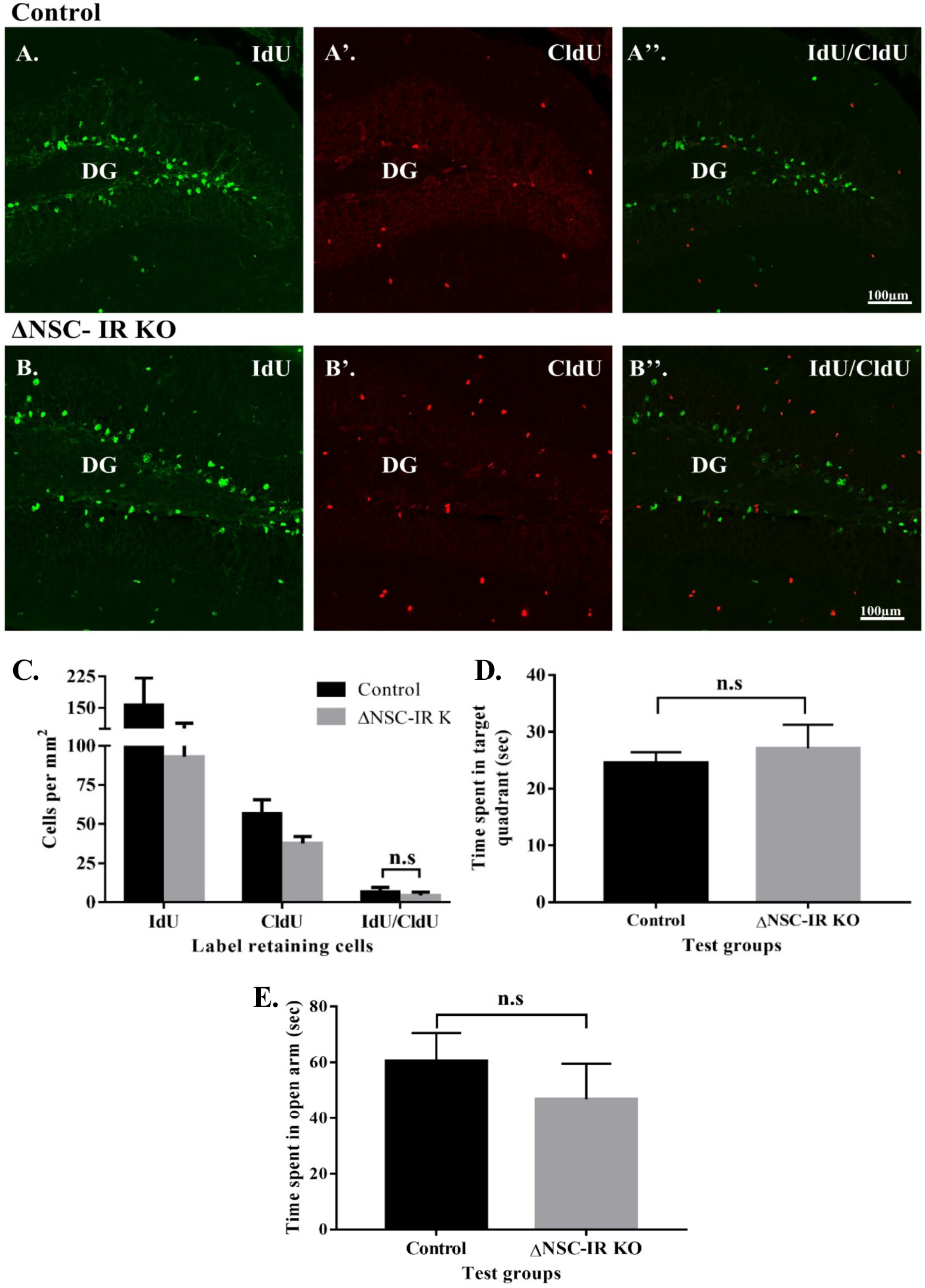
Hippocampal neurogenesis and hippocampal dependent behaviors are unchanged in ΔNSC-IR KO mice. **A.** Representative images of IdU and CldU labeling in control SGZ. **B.** Representative images of IdU and CldU labeling in ΔNSC-IR KO SGZ. **C.** Number of IdU, CldU single and double positive cells (Mean ± SEM, unpaired t test, not significant). **D.** Time spent in the target quadrant during the probe test of the Morris water maze (Mean ± SEM, unpaired t test, n=10 control, n=12 ΔNSC-IR KO). **E.** Time spent in the open arms of the elevated plus maze (Mean ± SEM, unpaired t test, n=10 control, n=12 ΔNSC-IR KO). Mice were analyzed 9 weeks after inducing IR deletion for label retention (Mean ± SEM, unpaired t test, n =4 control, n=5 ΔNSC-IR KO). Scale bar represents 100 µm.

**Supplemental figure 5.**
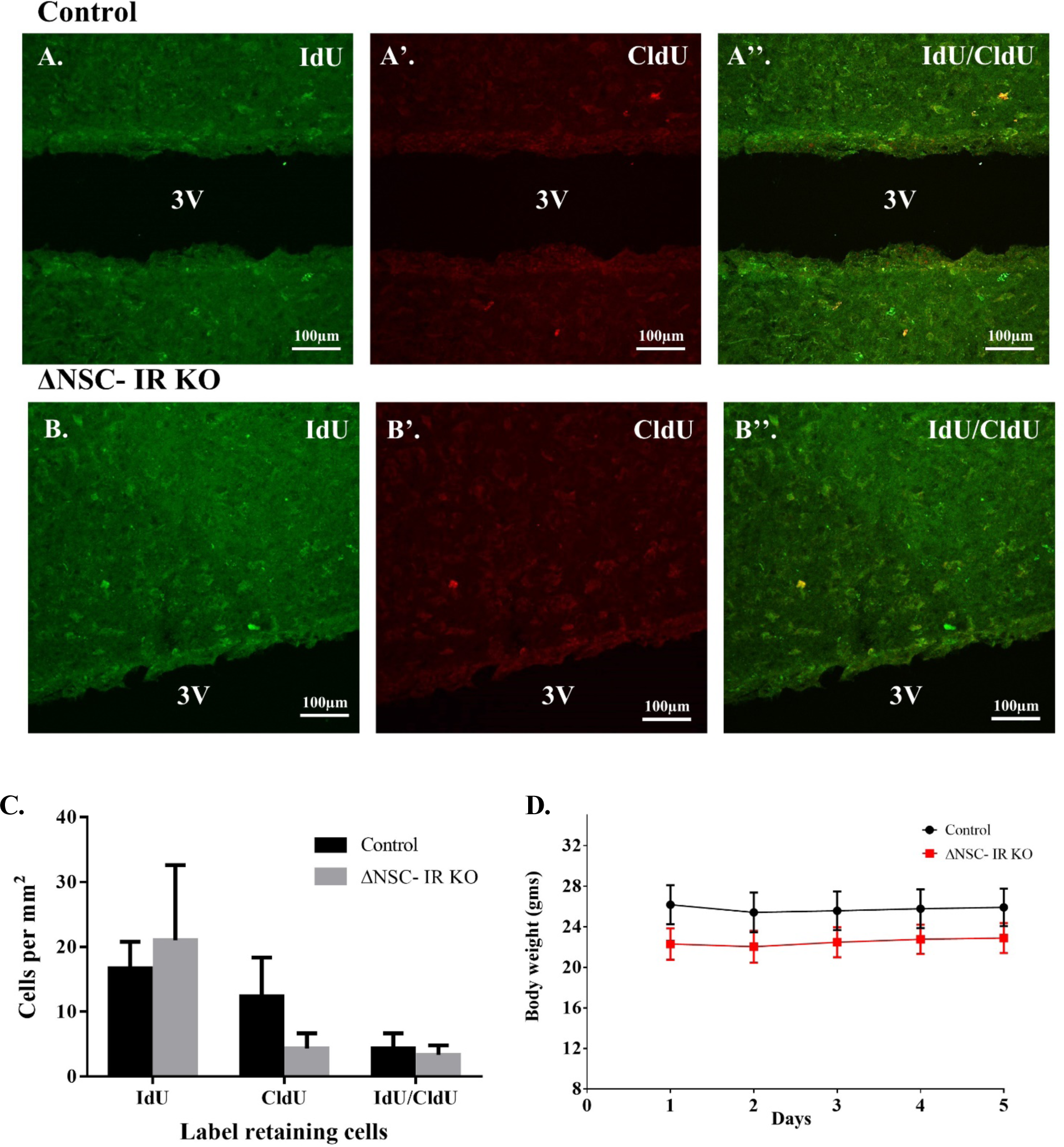
α-tanycytes in the 3^rd^ ventricle do not require IR signaling for self-renewal. **A-A’’.** Representative images of IdU and CldU labeling in control 3^rd^ ventricle (3V). **B-B’’.** Representative images of IdU and CldU labeling in IR KO 3^rd^ ventricle (3V). **C.** Number of IdU, CldU single and double positive cells in the median eminence (Mean ± SEM, unpaired t test, n=3 per genotype). **D.** Weight of control and IR KO mice over the 5 day injection period (Mean ± SEM, unpaired t test, n=10 control, n=12 IR KO). Scale bar represents 100 µm.

**Supplemental figure 6.**
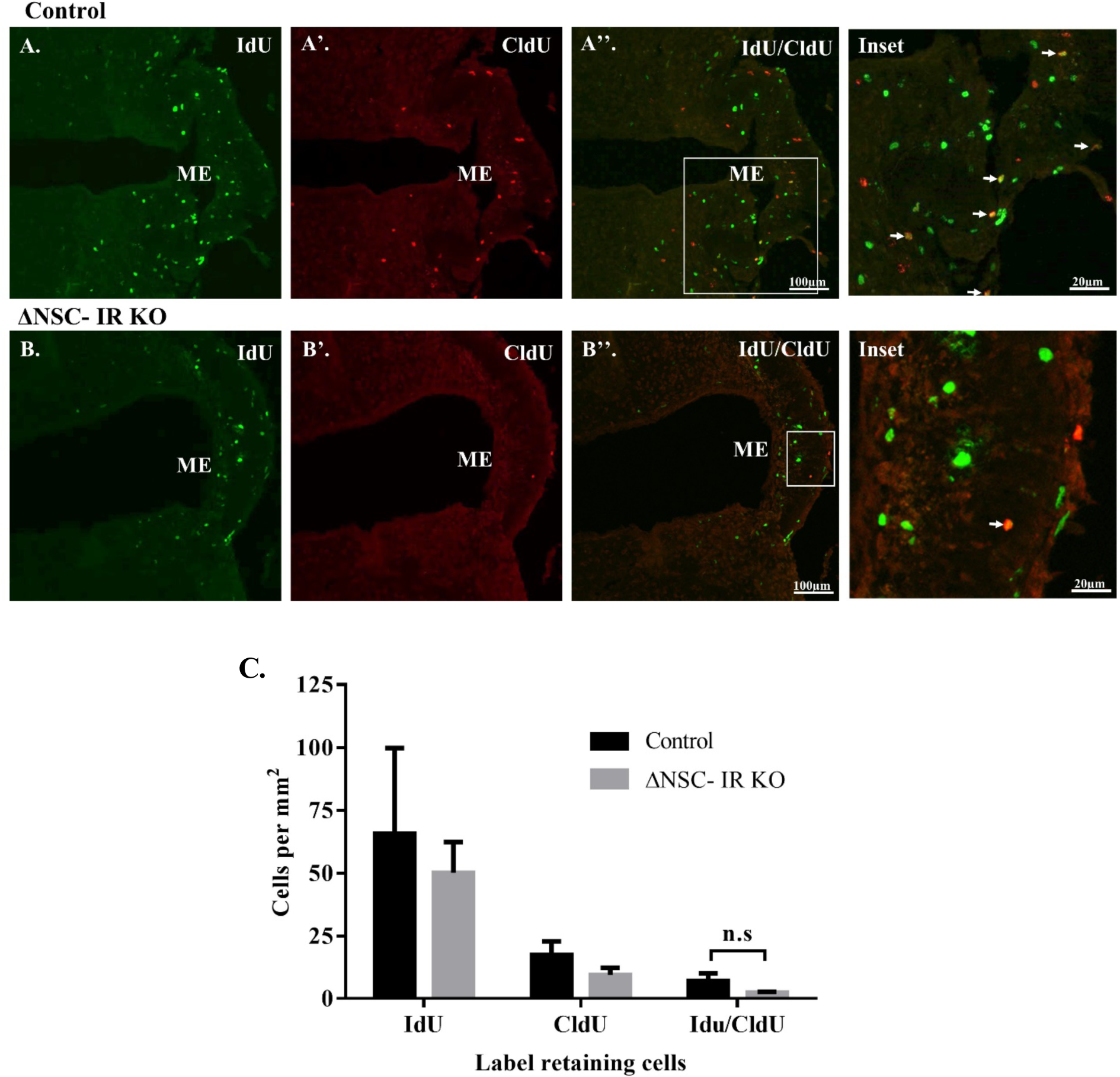
Hypothalamic median eminence may contain an IR responsive nestin+ sub-population of NSCs. **A-A’’.** Representative images of IdU and CldU labeling in control median eminence (ME). **B-B’’.** Representative images of IdU and CldU labeling in ΔNSC- IR KO ME. White arrows represent IdU^+^/CldU^+^ double positive cells. Scale bars represent 100 µm in panels A and B, and 20 µm for the insets. **C.** Number of IdU, CldU single and double positive cells in the ME (Mean ± SEM, unpaired t test, n=3 per genotype).

**Supplemental figure 7.**
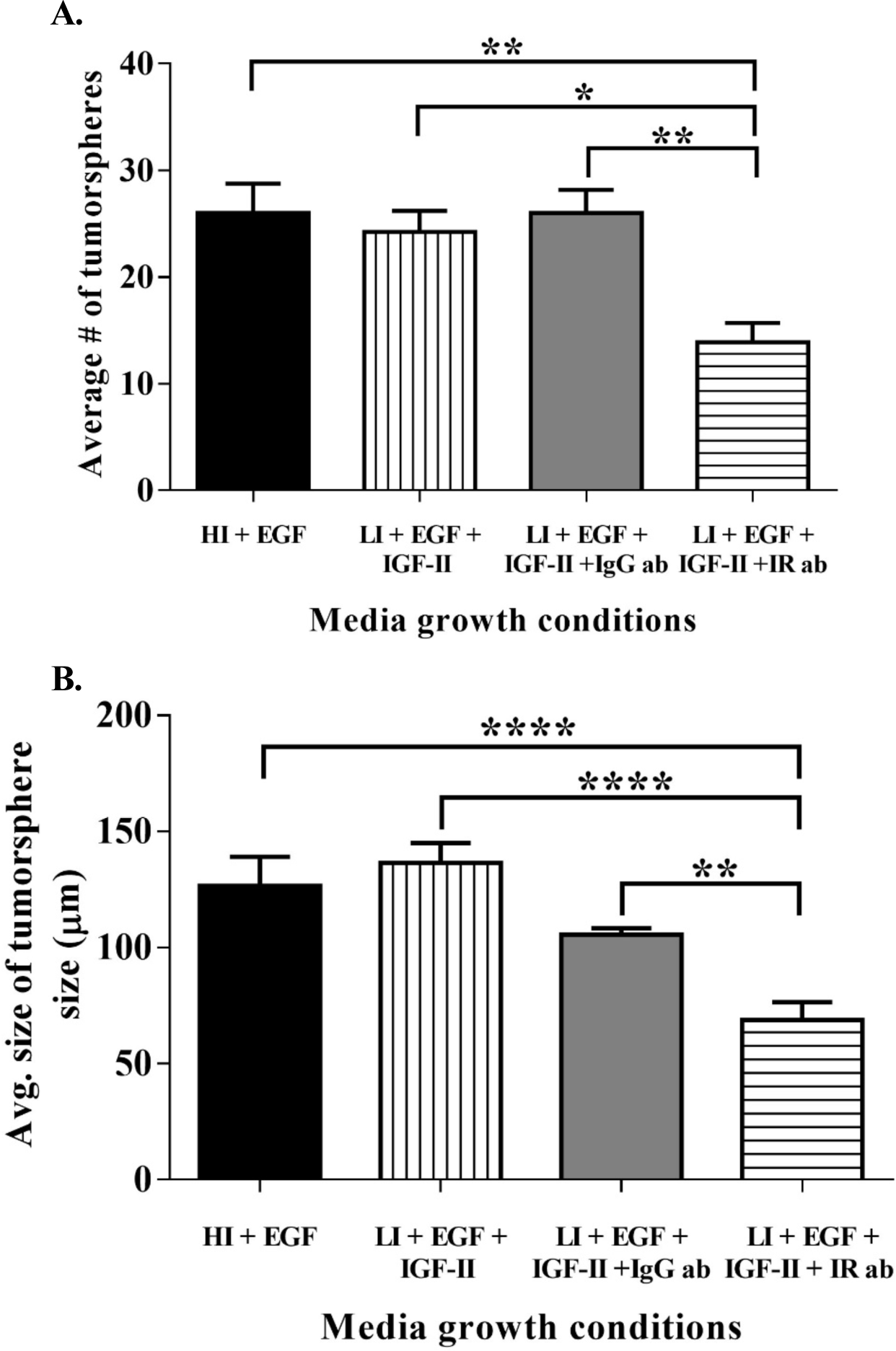
Mesenchymal tumorspheres grown with control antibody. **A.** Average number of tumorspheres grown in different media groups; HI+EGF vs LI+EGF+IGF-II+IR ab (**p<0.001), LI+EGF+IGF-II vs LI+EGF+IGF-II+IR ab (* p<0.01), LI+EGF+IGF-II+ IgG ab vs LI+EGF+IGF-II+IR ab (** p<0.001) **B.** Average size of tumorspheres grown in different media groups; HI+EGF vs LI+EGF+IGF-II+IR ab (*****p<0.0001), LI+EGF+IGF-II vs LI+EGF+IGF-II+IR ab (****p<0.0001), LI+EGF+IGF-II+ IgG ab vs LI+EGF+IGF-II+IR ab (**p<0.001). Data are from one experiment performed in triplicate. Mean ± SEM, One-way ANOVA.

## S. Chidambaram et al, Supplementary Methods

### Neurosphere propagation and quantification

Neurospheres were generated by enzymatically dissociating the periventricular region of IR^fl/fl^ pups (P4-5) as described previously {Ziegler, 2019 #13}. The cells were plated at a density of 2.5 x 10^5^ cells/ml in B27 media minus insulin supplemented with 20 ng/ml of recombinant human epidermal growth factor (EGF) (PeproTech) and 4.4 µM of insulin (Sigma) and passaged to secondary spheres (61). ImageJ software was used to measure neurosphere diameter, where any sphere that was at least 30 µm in diameter was defined as a neurosphere. Neurosphere number was calculated by counting 6 random fields per well and 3 wells per experiment.

### Flow cytometry

Spheres were dissociated by incubation in 0.2 Wünsch unit (WU)/ml of Liberase DH (Roche) and 250 μg of DNase1 (Sigma) in PGM solution (PBS with 1 mM MgCl2 and 0.6% dextrose) at 37°C for 5 min with gentle shaking. An equal volume of PGM was added and the spheres were placed onto a shaker (LabLine) at 225 rpms at 37°C for 15 min. After enzymatic digestion, Liberase DH was quenched with 10 ml of PGB (PBS without Mg^2+^ and Ca^2+^ with 0.6% dextrose and 2 mg/ml fraction V of BSA (Fisher Scientific, BP1600-100) and cells were collected by centrifugation for 5 min at 200 x g. Cells were dissociated by repeated trituration, collected by centrifugation, counted using ViCell (Beckman Coulter, Miami, FL) and diluted to at least 10^6^ cells per 50 μl of PGB. All staining was performed in 96 V-bottom plates using 150 μl/well. For surface marker analysis, cells were incubated in PGB for 25 min with antibodies against Lewis-X (1:20, LeX/CD15 FITC, MMA; BD Bioscience), CD133-APC (1:50,13A4; eBioscience), CD140a (1:400, APA5; BioLegend) and NG2 Chondroitin Sulfate Proteoglycan (1:50, AB5320; Millipore). Cells were washed with PGB by centrifugation at 278 x g. Goat anti-rabbit IgGAlexa Fluor 700 (1:100; Invitrogen) was used for NG2. Cells were then incubated in LIVE/DEAD fixable Violet (Invitrogen) for 20 min for dead cell exclusion. Cells were washed with PGB by centrifugation at 278 x g. They were fixed with 1% ultrapure formaldehyde (50000; Polysciences, Inc) for 20 min, collected by centrifugation for 9 min at 609 x g, resuspended in PBS w/o Mg^2+^ and Ca^2+^ and stored at 4°C for next day analysis. All sample data were collected on the BD LSR II (BD Biosciences Immunocytometry Systems). Matching isotype controls were used for all antibodies and gates were set based on these isotype controls. Post-acquisition analysis was performed using FlowJoX (Tree Star Inc, Ashland, OR). Among the live cells only tdtomato positive cells were analyzed.

### Thymidine analogs and Dual Thymidine Analog Detection

IdU and CldU detection was performed as described by Tuttle et. al. 2010 (62) with the following modifications: cryosections of forebrain SVZ and hippocampus (30 µm) were washed with 0.3% of triton-X-100 for 30 min at RT. Antigen retrieval was performed using Sodium citrate for 20 min at 96⁰C in a steamer. Tissue was blocked using 10% donkey serum for one hour at RT.

IdU (1:100) was added to tissue overnight at 4⁰C. The following day slides were washed with low salt TBST solution (36mM Tris, 50mM NaCl, 0.5% tween-20; pH 8.0) for 20 min at 37⁰C at 225 rpm. CldU antibody (1:100) was added overnight at 4⁰C. IdU was used with dylight 488, CldU with dylight 647 and DAPI at 1:5000 in PBS for 15 min. Slides were mounted in fluorogel (Electron Microscopy Sciences 17985).

### Morris Water Maze

*Hidden Platform*: The set up and testing for the water maze was as previously described {Ziegler, 2019 #13} (see also Supplementary Methods). Mice were tested in the Morris Water Maze for 5 consecutive days. Day 1 consisted of 1 pre-training session performed with the platform visible. For the pre-training session, mice were placed on the platform for 10 seconds, allowed to swim for 20 seconds, and then placed on the platform for another 10 seconds; each mouse received 4 trials. On days 2-5, mice received 2 training sessions per day. For each training trial, mice were placed in the maze and allowed 60 seconds to find the hidden platform, the latency to find the platform was measured for each trial. Each training session consisted of 6 trials with the mouse placed in a different quadrant of the maze each time, omitting placement in the TQ. On day 6, mice received a probe trial: the platform was removed, and mice were placed in the maze and allowed to explore for 60 seconds; the time spent in the TQ was measured.

### Elevated Plus Maze

Briefly, the animal was placed on the EPM apparatus facing the open arm. The time was recorded, and the animal was tracked using a video camera placed on top of the EPM. The animal was tested on the EPM for a total of 5 mins. After this the animal was removed and placed back into the cage. The AnyMaze software was used to track the total amount of time spent in the open arms, indicating less anxiety.

### Expression analysis of RNAseq data

RNA was obtained from cultures of stem cells isolated from two glioblastoma tumors (code-named GB2, WCR8) and from one normal human neural stem cell culture (HNSC). Next-generation RNA-sequencing was performed in the Epigenomics Core Facility of the Albert Einstein College of Medicine, NY, using an Illumina HiSeq2500 machine. Detailed protocols for library preparation can be found at http://wasp.einstein.yu.edu/index.php/Protocol:directional_WholeTranscript_seq. Sequence reads were aligned to the human genome (hg19 build) using *gsnap* Genomic locations of genes and exons, as defined in Refseq, were extracted from the *refGene.txt* file (http://hgdownload.cse.ucsc.edu/goldenPath/hg19/database/refGene.txt.gz). Read summarization at the gene level was done for all the genes in Refseq using the bam alignment files and in-house scripts, taking only reads with mapping quality of 20 or greater. The number of raw reads mapping to a gene was standardized to reads per kilobase per million reads (RPKM). The number of RPKM for the insulin receptor (INSR) and for the insulin-like growth factor 1 receptor (IGF1R) was extracted and the ratio of INSR to IGF1R was obtained for each sample.

